# Physarum polycephalum: Establishing an Assay for Testing Decision-making Under Shifting Somatic Boundaries

**DOI:** 10.1101/2021.10.17.464734

**Authors:** Samuel P. Levin, Michael Levin

## Abstract

Prior studies of decision-making generally assume a fixed agent which maximizes utility among its various options. Physarum polycephalum is a popular model for basal cognition that can be cut into pieces that may or may not re-join. We exploited this capacity to develop a novel assay in which radical changes to the agent itself are among the options of the decision-making process. Specifically, we transected a Physarum culture in the presence of a food reward that was located closer to the new smaller piece. In this scenario, the newly created branch must choose between exploiting the reward itself, or first re-connecting with the original mass (and sharing the nutrient reward across a large body). We report a pilot study establishing a protocol in which the number of agents is part of the decision-making process. We observed that despite the presence of food, new branches strongly prefer to merge back to the syncytium before exploiting the reward. Many improvements to the protocol are possible, to extend this effort to understand the interplay between behavioral options and the structure and boundary of the individual making choices in its environment.

## Introduction

Numerous fields of cognitive science, psychology, and economics focus on agents (rational or basal) making decisions to optimize their welfare. Most studies of decisionmaking behavior have in common one key feature: the agent is constant throughout the study: a single organism acts to maximize some sort of utility. We sought to expand this paradigm by establishing an assay in which the agent itself is plastic, and can undergo radical transformation as one of the possible actions. In this case, the calculus of comparing options becomes much more complex because the beneficiary of any reward is not necessarily the same individual that is making the decision. In one sense, this would be an empirical implementation of classic philosophical puzzles about personal identity, and the rationality of decisions made to benefit one’s future self, one’s clones, etc. More practically, we wanted to better understand the intersection between basal cognition (primitive decision-making) [1–3] and the plasticity of the biological process that gives rise to integrated Selves from living components [4–6].

The slime mold *Physarum* polycephalum is a popular model for studying problemsolving and basal cognition in aneural organisms [7]. Physarum is a rapidly-growing yellow slime mold, commonly found in forests. *Physarum* is perhaps most famous for the numerous experiments conducted on it by Nakagaki, Latty, and others, that revealed its talent for creating complex transportation networks and solving mazes [7–9]. Physarum is a simple and adaptable model for low-level biological decision-making [7, 10–19], but it also has an uncommon trait that makes it uniquely valuable: its anatomical plasticity and behavioral capacity under significant modification. Physarum is unicellular, but can grow very large and is an ideal organism for understanding how wholes made of parts make decisions.

Physarum can be cut in any location, producing multiple distinct entities which can later reconnect and once again become a single organism [20]. Thus, this organism’s “Self” is completely scalable and plastic, and the choice to rejoin the whole or become an independent organism is one that Physarum can make during any behavioral assay. Other organisms, especially neural ones, cannot be easily tested in this way. The slime mold system not only has the ability to determine whether or not it will share food with “others” it also has control over how many “others” there are altogether. Cutting a Physarum does not injure it, it just creates a second, equally competent Physarum. Thus, we designed a pilot assay to test Physarum’s behavior when the scale and number of Selves is as much an option for behavior as is the decision to exploit food resources.

In our assay, we cut a single existing Physarum into a large Physarum A and a small Physarum B, with or without a food reward being close to the new piece B. Physarum B would thus have two options: remain separate and collect the food for its new Self, or, rejoin A, and be once again part of a much larger mass, at the cost of having to share resources with the whole. If the new piece proceeds to exploit the food, it would suggest that a small piece rapidly acquires a new individuality and selfish behavior. On the other hand, if it first chooses to rejoin, then that would suggest that the Physarum’s strategy is to prioritize being part of a larger individual, and that a small piece does not rapidly acquire a self-protection behavior (is willing to merge its identity into a larger whole, in effect disappearing). This kind of calculus is interesting because the logic of selfish behavior is not straightforward: prior to a join, a selfish agent should want to keep the food to itself, but after the join, it will not exist, so there is no sense in which it can compete or share resources with “the other” (since it will be part of the other).

We hypothesized that after cutting, the small piece would proceed to exploit the food. Our preliminary results suggest the opposite, with pieces generally preferring to merge at the same rate and timeframe regardless of whether food resources were nearby.

## Methods

### Culture and Propagation

For our experiments, we used the LU352 strain provided by Audrey Dussutour, which we perpetuated as follows. All Physarum cultures were stored in a computer-controlled humidified habitat constructed from an Amazon insulated grocery box (Figure 1). Humidity and temperature were measured and controlled by an Inkbird Temperature and Humidity controller model ITC-608T. The humidity level was set to 85%. To determine the optimal humidity level, plates were placed in the habitat overnight with a small amount of Physarum in the center, with different humidity settings. The setting that saw the most growth was 85%, so we retained that value for future experiments. For humidification, we used a Coospider Mister 3L model humidifier. The Physarum was grown on Petri dishes filled with 500 grams of Fisher Bioreagents BP1423-500 Agar. The *Physarum* colonies were fed with Quaker brand 100% whole grain Old-Fashioned oats.

**Figure 1:**
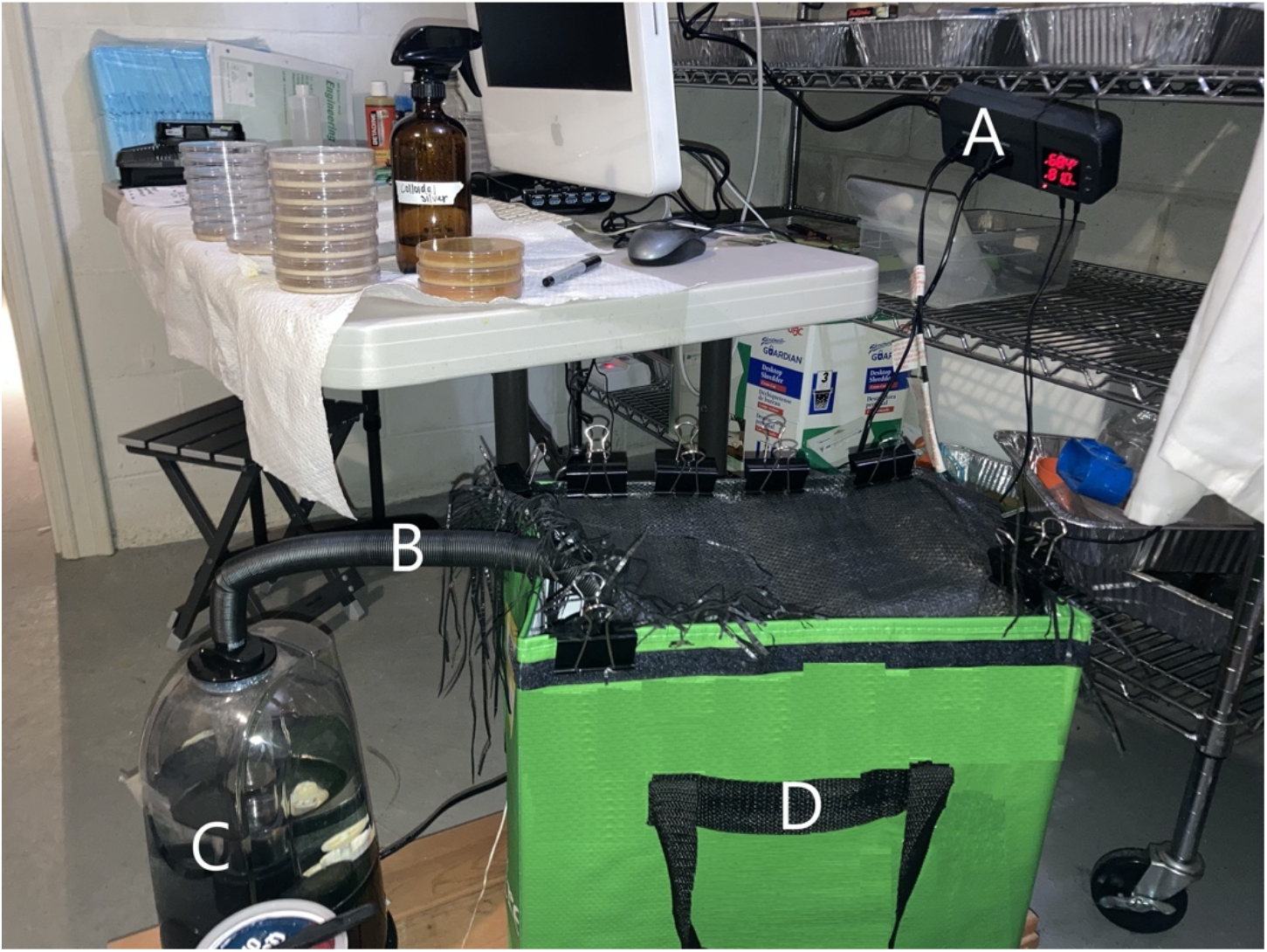
Culture apparatus. Shown here are the humidity controller (A), the humidifier pipe (B), the humidifier itself (C), and the habitat (D), with green section edited to remove irrelevant writing on the container).

The dishes were separated into sorted columns within the habitat, and with an additional column for stock plates. Stock plates (Figure 2) were plates where *Physarum* was allowed to grow unchecked with additional food frequently provided. Stock plates were not used directly in any experiments. When new colonies were necessary for experiments, sections were lifted out of these plates using a Corning Inc. model 3008 scraper and placed the chunk of agar with the *Physarum* facing down in the new plate.

**Figure 2.**
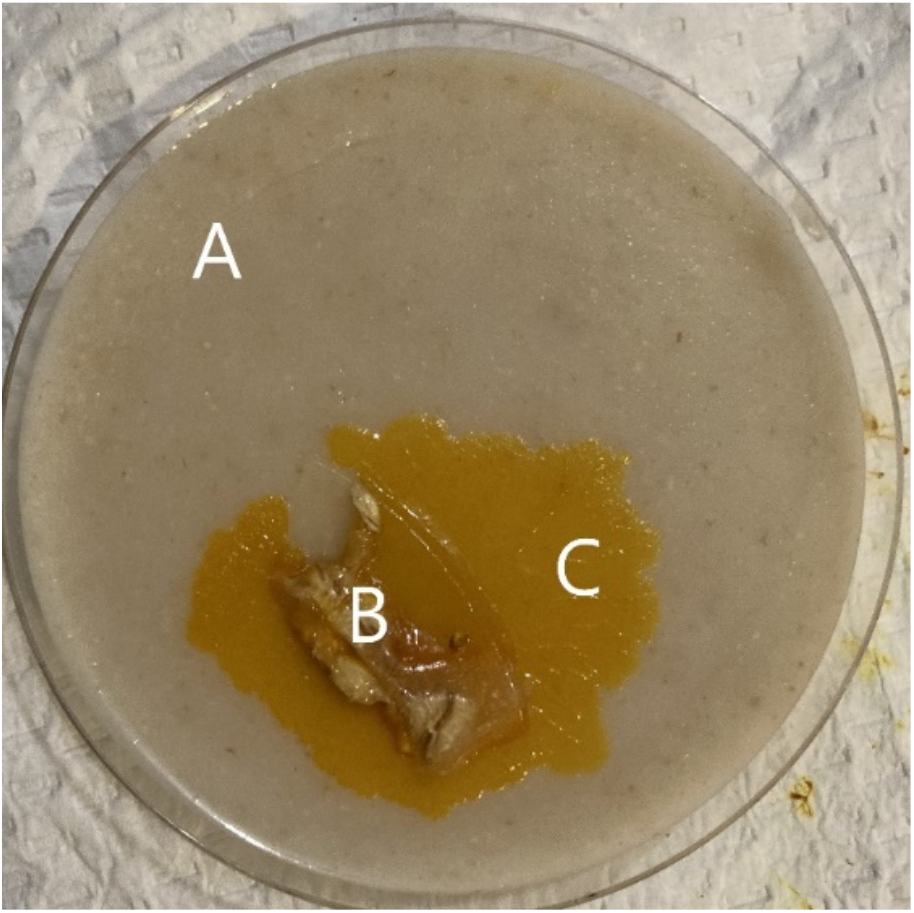
A Physarum stock plate. An example stock plate with Physarum in the early stages of its growth. A is the oat-saturated Agar. B is a transplant fragment of Physarum from a prior plate. C is newly grown Physarum.

**Figure 3.**
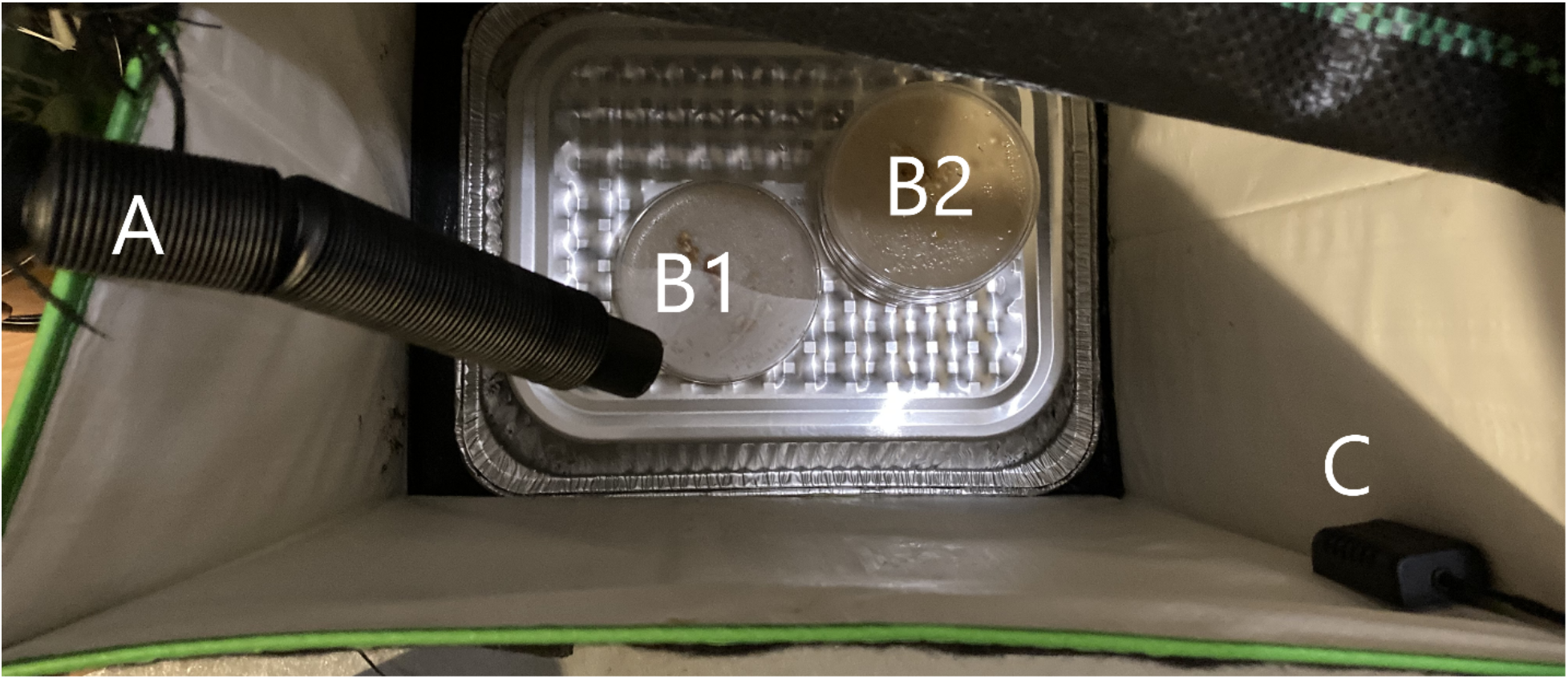
Physarum culture apparatus interior (with lid removed) A view inside the habitat. Pictured are the humidity input pipe (A), two stacks of Physarum plates (B1, B2), and the humidity censor (C).

To establish a baseline growth rate, we initially tested the time required for a Physarum to grow from one edge of a standard Petri dish to an oat on the other side. Once the dishes were prepared, we observed them hourly throughout the day. It was determined that the Physarum takes approximately 12 hours to cross a standard 100mm diameter plastic non-coated Petri dish. This measurement was used as a guideline for all following experiments, plates were set in the evening, and checked on in the following morning, approximately 12 hours apart.

## Results

### Establishing conditions that facilitate merging

The first round of testing involved checking the minimum viable distance for the Physarum to merge. Nine plates were set, divided into groups of 3. Each group had a different size of gap, “Small”, “Medium”, or “Large”. “Large” gaps had a width of ~2.28mm. “Medium” gaps were ~1.14mm wide, and “Small” gaps were ~0.5mm wide.

The gaps were created by lifting two (18.5mm x 13.4mm) sections of the Physarum out of an existing colony and placing them at a predetermined distance from one another. All nine plates died and failed to merge. This was determined to be caused by a cliff-like gap in the agar. A gap of 1.14mm or wider was large enough that the pressure of the agar would not cause it to push the gap closed and created vertical walls on either side of the gap, which the Physarum was not able to climb. A failure to merge is shown in Fig. 4.

**Figure 4.**
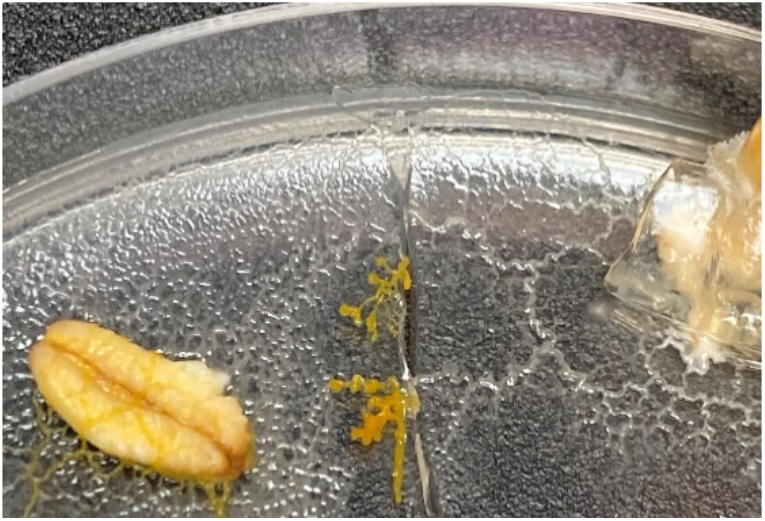
An unsuccessful Physarum merge. Example of an unsuccessful merge. Because there was no merge, the right side withered and died, lacking the nutrients the left side had access to.

When the same tests were repeated but with the cuts in the Physarum being made by simply lifting out a section of the agar, with nothing but the plastic of the Petri dish between them, the Physarum was able to merge at a “Small” distance easily (Fig. 5).

**Figure 5.**
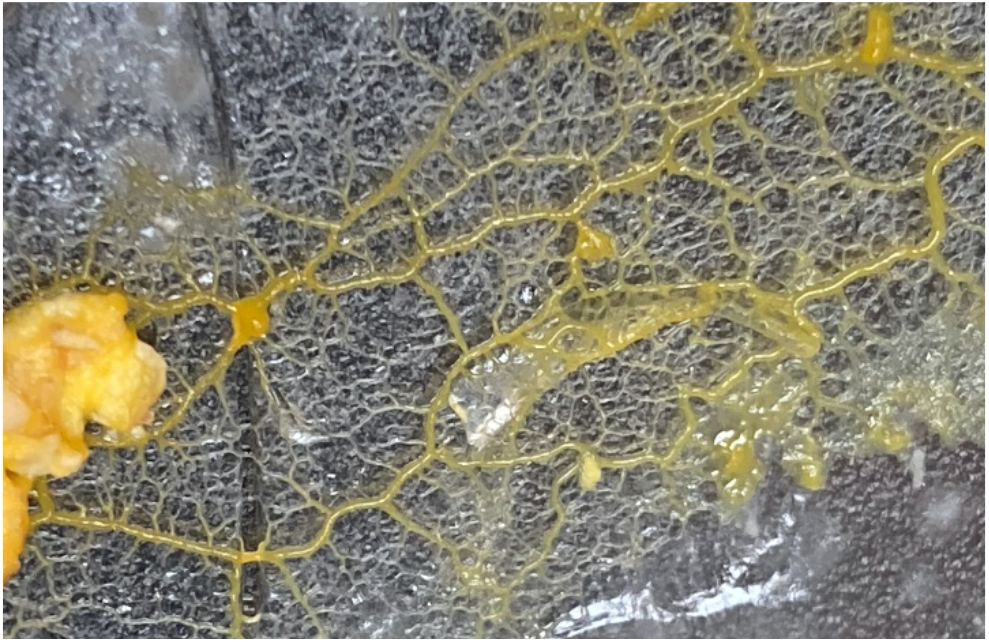
A successful Physarum merge. Example of a successful merge across a cut.

### Determining exploit vs. merge preferences

To determine if the Physarum would stay separate or cooperate, we first allowed the Physarum to colonize half of a Petri dish. One oat was placed in the dish prior to adding the Physarum, to encourage growth and keep it from starving while it established itself in the dish. 12 hours later, we used the blade of a Corning 3008 scraper to cut a straight line through the Physarum (the cut was approximately 0.5mm wide before the blade was removed and the agar pushed the cut closed).

After making the cut, a new oat was placed in the primary dishes, alongside the far wall, opposite to the Physarum. In the control dishes, the cut was made as described above, but no oat was added after cutting. The dishes were collected, and the results were observed after another 12 hours had passed. This set of procedures was repeated three times in phases separated by seven days, to allow the stock plates to replenish. There were 10 control and 10 primary dishes in each phase.

In each phase, 20 plates were established, separated into a group of 10 controls and 10 experimental plates. The combined results of the three experimental repeats are tabulated below. The experimental plates had an oat added after the cut was made, the control plates did not. Undetermined plates were plates where it was impossible to determine whether or not a merge had occurred with human eyes or a camera. The results are displayed in Table 1.

**Table 1:** Results of merging tests

|  | **Experimental plates**<br>**(with oat)** | **Control**<br>**(no oat present)** |
| --- | --- | --- |
| Merge | 22 | 23 |
| No Merge | 5 | 4 |
| Undetermined | 3 | 3 |

A Chi-Square test on these data (excluding undetermined values) showed that the samples were not significantly different (0.715). Since the data do not enable us to reject the null hypothesis (that behavior with and without oat are indistinguishable), we conclude that within our study, the new Physarum piece preferred to merge rather than exploit the food.

## Discussion

We set up an assay in which a newly-created individual had the opportunity to merge back into its parent body or exploit a resource resulting in a much higher density of food per biomass. Within the limits of our study, we observed that the Physarum appears to exhibit a strong preference for merging, regardless of the presence or absence of an oat reward. This suggests the Physarum prefers the long-term benefit of being a single-large entity over the short-term benefit of monopolizing a single item of food. It is plausible that ecologically, the observed strategy is the more adaptive one and has been selected by evolution. It is also possible that the Physarum could be remembering being a single entity and be motivated to return to that state as a kind of homeostasis. Physarum is known to be capable of remembering past experiences [21–24], and it is possible that the rapid merging is a kind of regenerative/repair response (anatomical homeostasis) that acts faster than individual pieces can acquire an independent identity.

### Limitations of Study

There were a number of limitations to this study, which we view as a first step of establishing a novel assay to address issues at the intersection of basal decision-making and somatic scaling. These experiments will be repeated with larger numbers, and precise testing of different distances of gap, oat position, etc.

One possibility is that something about the construction of the experiment was motivating the *Physarum* to merge as quickly as possible. The most obvious potential trigger for such behavior would be the distance between the back half of the cut Physarum and the oat. If the energy cost of merging is lower than the cost of traveling far enough to reach the oat before the front half does, perhaps merges immediately to save time and resources.

Another limitation was that this study relies on qualitative, visual observations by a human observer. The Physarum “moves” by growing towards objects of interest and letting the sections behind it wither away (these sections rapidly turn translucent and almost invisible). When the Physarum is cut and the oat placed in the dish, it (both the cut and uncut segments) starts “moving” towards the oat. This means that on many occasions (10% of the tests, as previously mentioned), even before the merge had/had not occurred, the Physarum around the area of the cut had already begun turning transparent. By the time the Physarum had entered the merge window, it was already completely transparent around the cut area, and impossible to tell what had occurred.

### Future Work

Future tests will be conducted by testing whether or not the Physarum had merged in a way unaffected by an observer’s limitations – an objective quantification, ideally using automated image processing techniques. One possible method to achieve this would be to dye the oat added to the plates a distinct color so that when the Physarum transferred the nutrients from the oat to itself, the dye would flow with it and discolor the whole mass. This would prevent older sections from becoming invisible, and make observation quicker and more precise.

The next tests should also seek to make the conditions as equal as possible for both halves of the cut Physarum. While not necessarily possible in the cramped spaces of a small Petri dish, the next set of experiments would ideally place the two chunks of Physarum with equal distances between each other and the oat(s). This would prevent the issue of travel distance potentially biasing the Physarum’s decision-making.

The Physarum’s convenient nature and flexibility make it an ideal model system to address the many fascinating issues surrounding scale of self and basal cognition. In addition to testing different Physarum strains under different conditions, it will be interesting to place a temporary barrier along the cut for different periods of time to determine whether there is a time period at which the small piece acquires individuality and begins to exploit the food without merging.

### Broader implications

These pilot experiments suggest a whole category of assays in similar types of model systems to develop for modeling decision-making in which the selfish Self can change during the experiment – a sort of second-order system in which maximizing reward is a dynamic process because the beneficiary of the reward is changing in realtime as a result of decisions and behavior. This will apply naturally to work in swarm intelligence and collective decision-making, both biological and robotic [25–33]. But it is important to note that other model systems do also enable the study of cognition during radical change of the organism, including the caterpillar-butterfly transition [34, 35] and the retention of memory in planaria that have to regenerate entire brains [36, 37].

The most obvious application of these ideas is to biological scenarios where competent agents scale up to form new levels of agency, such as cells which make bodies. In cancer, cells revert to an ancient unicellular lifestyle, abandoning their collective morphogenetic goals and treating the rest of the body as external environment [4, 38, 39]. While numerous studies of cancer from the perspective of game theory have been made [30, 40–46], further process could be achieved by better understanding this process not from the perspective of increased selfishness, but constant selfishness coupled with changes of the size of the Self, and thus changing incentives.

There are also many parallels that could eventually be forged between such experiments with societies of cells and subcellular components (as in Physarum, which is all one cell), and altruistic behavior in psychology. Much work has been done on individual human (and non-human animals’) decisions to share resources with the larger group. Interestingly, while not highly amenable to experimental manipulation, even human bodies are relevant to division of Selves below the society level. These include brain hemisphere dissociation (and someday, transplant) scenarios [47, 48], and multiple personality disorders in which the number and identity of the beneficiary of therapy is changing during the process (e.g., integration therapy that literally dissolves one or more human personalities) [49–51].

There are many interesting philosophical questions that are raised about rational decision-making when the rational agent itself may not be there to benefit from them. On a more practical level, understanding how biology handles adaptive behavior in the context of scaling higher-order agents from potentially competent parts will shed light on many aspects of evolution, cell biology, and cognitive science.

## Acknowledgements

We thank Jayati Mandal for plate preparation, and Audrey Dussutour for providing the LU352 strain Physarum used for this study.

